# WEBNG: A templating tool for weighted ensemble sampling of rule-based models

**DOI:** 10.1101/2022.10.28.514312

**Authors:** Ali Sinan Saglam, James R. Faeder

## Abstract

Time scales for biological processes span many orders of magnitude, forcing modelers to tackle coupled processes that have large time scale gaps. This results in rare events, which take longer to occur than the fastest processes in the model. Efficient generation of rare events has been a focus of modelers for a long time and multiple software packages implement various rare event sampling algorithms. However, these packages frequently require expertise to get started with, making it harder for researchers to start using them. WEBNG (short for Weighted Ensemble–BioNetGen) is an open source software framwework that bridges the open source software packages WESTPA, which implements the weighted ensemble method for sampling rare events, and BioNetGen, which facilitates the specification and simulation of biochemical reaction network models following a rule-based approach. WEBNG simplifies rare event sampling in simulations of rule-based models by taking a model specified in the BioNetGen language (BNGL) and generating a WESTPA simulation folder ready to simulate with default parameters selected to match model observables. WEBNG is written in Python with dependencies only on proven, open-source packages that are in active development, which makes WEBNG easy to install and maintain. Here, we describe the architecture and features of WEBNG and demonstrate its capabilities through application to a two-gene model of cell fate transitions.

## Introduction

Biological molecules can undergo many transformations, including phosphorylation, binding, ubiquitination, etc. The multisite/multivalent nature of many molecular components of biological networks gives rise to combinatorial complexity in the number of individual species (complexes) and reactions that can occur in the reaction network governing system dynamics [1]. To simplify modeling such systems, rule-based modeling approaches were developed to avoid manually enumeration of the network [2,3]. Rule-based models define rules that specify how species can interact, and these rules are the applied iteratively to generate the network [4]. BioNetGen (BNG) [3,5,6] is an open-source rule-based modeling software package that allows for model construction in BioNetGen language (BNGL) and simulation of BNGL models.

Rare events in modeling are events that take a long time to occur compared to the time step used to simulate the model. Time scales on which biological processes happen span many orders of magnitude. Most biological models need to tackle coupled processes that happen at drastically different time scales [7], resulting in rare events. While in some models it is possible to simplify some of the faster processes to close the gap between time scales, in many models rare events are unavoidable because the fastest process in the system must be modeled for accuracy. Models with rare events can be computationally expensive to simulate because it takes many events to get a statistically robust estimate of any aspect of a biological process, which means the model has to be simulated for a long time in order to sample enough rare events.

Tackling this problem has been a focus of the computational modeling field and many rare event sampling methods have been developed [8–12]. One of these methods is called weighted ensemble path sampling method [8]. Weighted ensemble path sampling works by organizing multiple parallel trajectories and resampling them at a fixed time interval to replicate trajectories that are making progress towards the rare event of interest and terminate trajectories that are not. Each trajectory is assigned a statistical weight that is tracked and updated during resampling in order to ensure that each trajectory that is generated has the correct statistical weight for computing system properties.

WESTPA is an open-source, scalable, and interoperable software package that applies the WE strategy [13]. WESTPA has been successfully used to study multiple challenging biological processes [14–18] and is being actively developed. The fact that WE does not bias the kinetics of the model makes it easy to use with any model simulated with stochastic dynamics, including rule-based models. WESTPA has been designed with a general interface for stochastic simulation engines making it straightforward to couple with BioNetGen’s ssa method, which implements an efficient version Gillespie’s Direct Method [19] for stochastic simulation of reaction networks [6].

To this end we have developed a tool called WEBNG (Weighted Ensemble-BioNetGen), which generates a WESTPA template for rare event simulation of a BNGL model. The user provides basic parameters for a WESTPA simulation along with a BNGL model to simulate, and the tool creates a WESTPA simulation folder that can then be run like any other WESTPA simulation. WEBNG is designed to tackle high dimensional WE simulations that are common for rule-based models and also provides several built-in analyses that are tailored to network-level simulations (as opposed to the molecular level simulations for which WESTPA was originally designed). WEBNG aims to lower the entry barrier for systems biology researchers to use the weighted ensemble method to explore models where rare events occur in an efficient way without having to learn details of the WESTPA interface that are needed for advanced applications.

## Results

### WEBNG Architecture

WEBNG comes with three subcommands that will set up the WESTPA simulation folder and analyze the resulting simulation in steps (Fig. 1). The first subcommand, template, allows the researcher to point to the BNGL model file they want to simulate and this subcommand will generate a YAML file that is easy to read and modify and is populated with reasonable default values for package locations, WESTPA simulation parameters, and parameters for the built-in analyses, all of which can be modified by the user.

**Figure 1.**
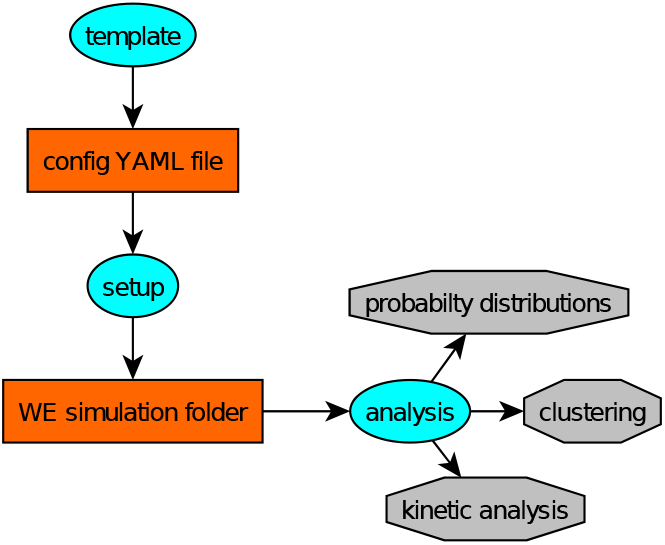
General workflow for WEBNG. Elliptical nodes are WEBNG subcommands, rectangular nodes are files that are generated and passed between subcommands, hexagonal nodes are analysis results.

The next subcommand, setup, takes as a single argument the name of the YAML file generated in the previous step. This subcommand will use this information to generate the standard WESTPA simulation folder, which contains all of the files needed to carry out a simuluation using WESTPA. From this point the user can add more sophisticated options that are not currently supported by WEBNG prior to invoking WESTPA. For information on what options WEBNG currently supports, see the WEBNG documentation linked at the end of this paper. This workflow allows a new user to quickly start simulating their

BNG model with reasonable defaults while also allowing an advanced WESTPA user to change simulation parameters and options freely. One important thing to note, by default WEBNG uses an adaptive Voronoi binning scheme [20] for the progress coordinates, which is a good starting point for high dimensional problems in the absence of further information. Binning strategies will be further discussed in the Results section.

Once the simulation is run, the user can use the analysis subcommand, which also takes in the same YAML file generated in the template step. This subcommand will take in the parameters provided in the YAML file and run the enabled analyses on the simulation. The built-in analyses shown in Fig. 1 are probability distributions, clustering, and kinetic analysis. Here, we provide brief descriptions of each analysis method with detailed examples provided in the next section.

There are two built-in probability distribution analyses, one that shows the joint probability distribution between every progress coordinate on a heatmap, and one that shows the evolution of the probability distributions of each progress coordinate. The joint probability distribution analysis is particularly useful to assess binning efficiency and simulation convergence.

WEBNG allows for automated clustering using the Generalized Robust Perron Cluster Cluster Analysis (GPCCA+) [21] implemented in the PyGPCCA+ python library [22]. GPCCA+ is a generalized version of the clustering algorithm PCCA+ [23,24], which is a clustering method based on the kinetics of a system. PCCA+ was originally designed to work on reversible systems under equilibrium, GPCCA+ extends this method to handle non-reversible systems as well. The user provides the number of clusters they want, and the GPCCA+ algorithm finds the ideal clustering of the bins in which transitions between clusters are minimized and transitions within clusters are maximized, maximizing the stability of each cluster. The resulting clustering can also be used in conjunction with WESTPA tools to calculate rate constants between clusters. Finally, WEBNG uses the clusters determined by GPCCA+ and the calculated transition matrices to make a GraphML file [25]. This file can be opened in Gephi [26] to visualize the clusters and transitions between them.

### WEBNG Installation

WEBNG is distributed via the Python Package Index (PyPI), which simplifies installation to a single command from the command line. Installing WEBNG also installs PyBioNetGen (a Python package that comes with BioNetGen) and WESTPA. This not only allows the user to avoid having to install the other two packages separately but also enables the template command to automatically locate the other packages and provide their paths in the YAML configuration file.

### Application of WEBNG to a Gene Expression Model

Tse et al. used weighted ensemble to run stochastic simulations of multiple gene regulatory network models, identify different phenotypes of the model, and estimate mean first passage times between those phenotypes. We used this study as a guide in designing of WEBNG with the aim to quickly set up simulations for the systems used in the paper and replicate the analyses in a streamlined fashion. Here, we have replicated the results of the exclusive mutual inhibition and self-activation (ExMISA) model shown in Fig. 2. In this model, two genes, A and B, encode transcription factors that activate their own transcription and repress the transcription of the other gene. The model includes the creation and degradation of the transcription factors as well as binding and unbinding of these transcription factors to the regulatory regions of both genes.

**Figure 2.**
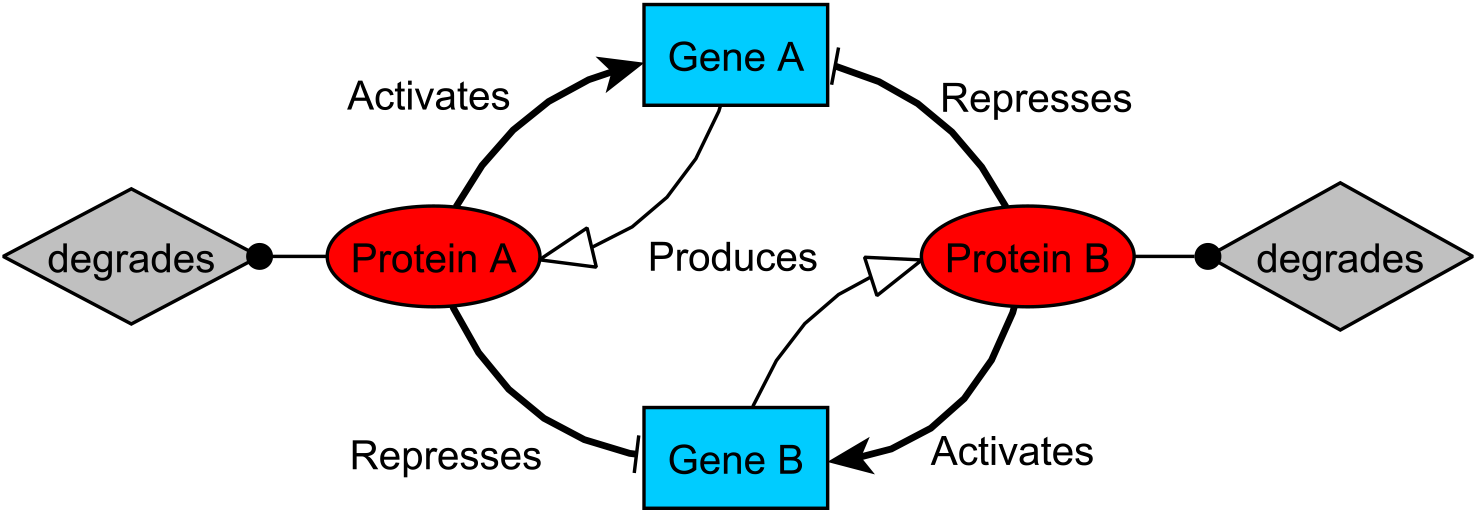
Diagram of the genetic switch model ExMISA [27].

The analysis reveals four expected high probability regions (Fig. 3(A)), one region where both proteins are low in count (lo/lo), one where both proteins are high in count (hi/hi) and two regions where one protein is low and the other is high in count (hi/lo and lo/hi). The automated clustering correctly identifies each cluster (Fig. 3(B)) and automated network visualization can visualize each state and transitions between them (Fig. 3(C)). Standard WESTPA tools can also be used to calculate the rate constants and therefore mean first passage times (MFPTs) between phenotypes identified from clustering (Fig. 3(C)).

**Figure 3.**
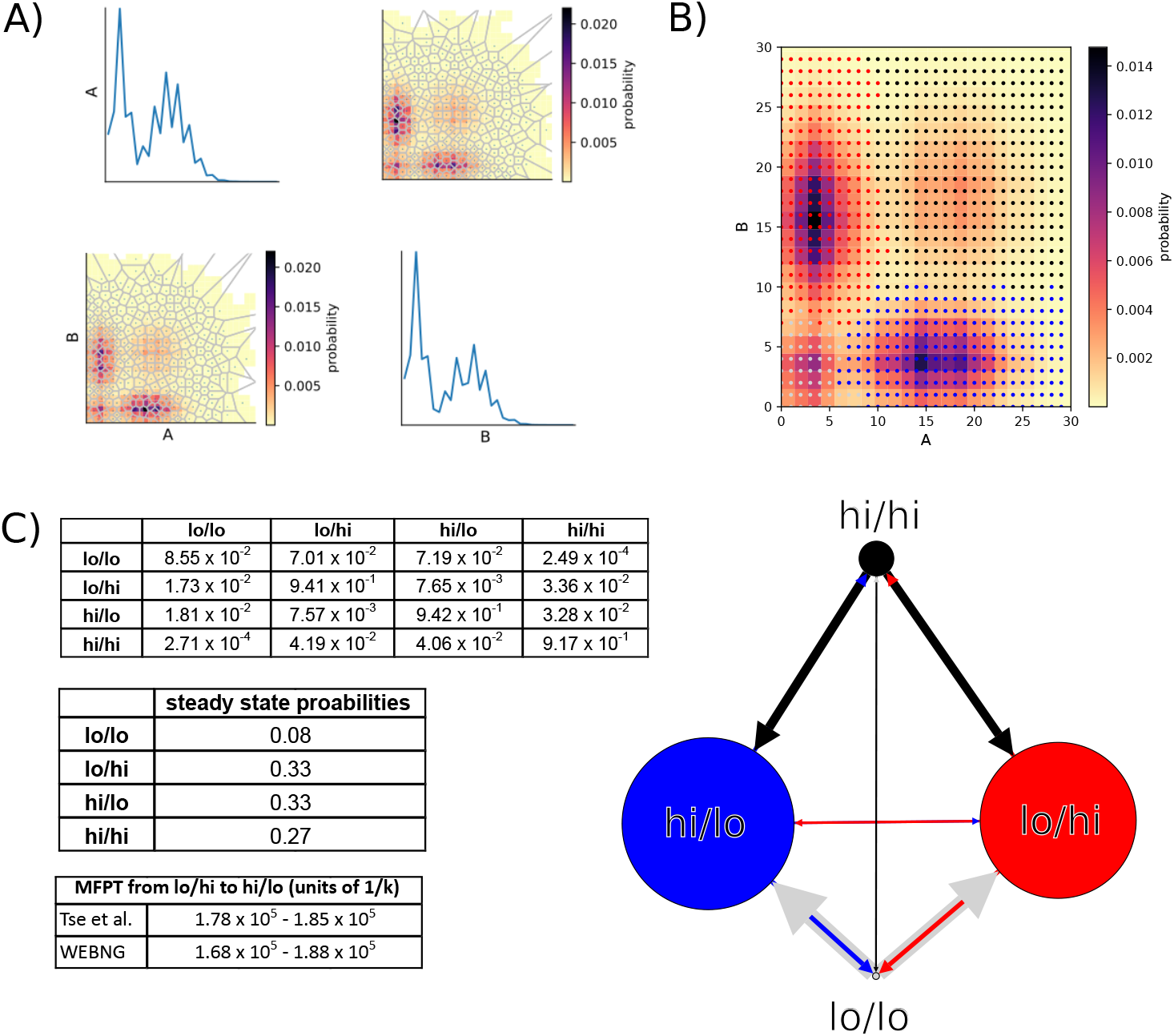
Diagram of the genetic switch model ExMISA [27] and results of automated analyses on a WE simulation. **A-C**, The WE simulation was run for 500 iterations (A) 2D heatmaps of joint probability distributions of protein A versus protein B counts. Lighter colors show less probable regions while darker colors show more probable regions. The probability distributions are estimated over the last 100 WE iterations. The grey lines indicate Voronoi bins used in adaptive binning. (B) Automated clustering correctly identifies each high probability phenotype. Clusters are represented by colored dots overlaid on top of a 2D heatmap of joint probability distribution of protein A counts versus protein B counts. Each colored dot represents one discrete possible state the model is in and each color represents a different cluster. (C) Transition probabilities between GPCCA+ generated clusters and steady state probabilities estimated from those transitions. MFPT is estimated using WESTPA tools and using definitions that match Tse et al. study. Network graph is automatically generated from the transition matrix and the steady state probabilities. Thicker edges mean higher probability transitions between clusters and node sizes scale with the probability of each phenotype in steady state. Each edge is colored by the state that they start from.

## Discussion

WEBNG is a simple templating tool that bridges two open software packages, WESTPA and BioNetGen, and lowers the barrier to entry to weighted ensemble rare event sampling of rule-based models. WESTPA, BNG and WEBNG are all well documented, which helps any potential researcher who wants to model processes that contain rare events using rule-based modeling by providing a simple starting point. BioNetGen also has the capability to take as input any model encoded using the Systems Biology Markup Language (SBML) standard [28]. Using PyPI as the distribution point not only makes installation simple but also allows for all three software packages to be easily kept up-to-date and in sync. New analyses, higher simulation efficiency, and increased support for WESTPA parameters are currently in development for WEBNG.

Another advantage of a dedicated software package for templating these simulations is reproducibility. There are many studies where the model is implemented in a language-specific manner (including the Tse et al. [27], in which the WE simulation is coded in MATLAB) and contain hard-coded variables which makes reproducing results challenging even when the code is provided with the published paper. Dedicated software packages like WEBNG allow for the same simulation setup to be generated as long as the original model file is provided and results analyzed in a consistent manner to make results more reproducible.

More information on BioNetGen, WESTPA, and WEBNG can be found at the following links: https://bionetgen.org/, https://westpa.github.io/westpa/ and https://webng.readthedocs.io/en/latest/.

If you encounter any problems using WEBNG please report your problem under the GitHub issues page https://github.com/ASinanSaglam/webng/issues.

## Acknowledgments

We thank Elizabeth Read for helpful conversations about the two-gene model, Caleb Armstrong for proofreading and feedback on the manuscript, and Philipp Schlegel for the LATEXtemplate, which can be found at https://www.overleaf.com/latex/templates/arxiv-slash-biorxiv-template/phncddwqtxpc. We gratefully acknowledge funding from NIH grants P41 GM103712 and R01 GM115805.

## Notes

### Competing Interest Statement

The authors have declared no competing interest.

### Summary of Updates

Updated the description and references for the clustering algorithm used by WEBNG.

## References

1. Hlavacek WS, Faeder JR, Blinov ML, Perelson AS, Goldstein B. The complexity of complexes in signal transduction. Biotechnology and Bioengineering. 2003;84(7):783–94. doi:10.1002/bit.10842.

2. Boutillier P, Maasha M, Li X, Medina-Abarca HF, Krivine J, Feret J, et al. The Kappa platform for rule-based modeling. Bioinformatics. 2018;34(13):i583–i592. doi:10.1093/bioinformatics/bty272.

3. Faeder JR, Blinov ML, Hlavacek WS. Rule-based modeling of biochemical systems with BioNetGen. In: Systems Biology. Springer; 2009. p. 113–167.

4. Blinov ML, Yang J, Faeder JR, Hlavacek WS. Graph theory for rule-based modeling of biochemical networks. In: Transactions on Computational Systems Biology VII. Springer; 2006. p. 89–106.

5. Blinov ML, Faeder JR, Goldstein B, Hlavacek WS. BioNetGen: software for rule-based modeling of signal transduction based on the interactions of molecular domains. Bioinformatics. 2004;20(17):3289–3291.

6. Harris LA, Hogg JS, Tapia JJ, Sekar JA, Gupta S, Korsunsky I, et al. BioNetGen 2.2: advances in rule-based modeling. Bioinformatics. 2016;32(21):3366–3368.

7. Peters B. Reaction rate theory and rare events. Elsevier; 2017.

8. Huber GA, Kim S. Weighted-ensemble Brownian dynamics simulations for protein association reactions. Biophys J. 1996;70(1):97–110. doi:10.1016/s0006-3495(96)79552-8.

9. Dellago C, Bolhuis PG, Chandler D. Efficient transition path sampling: Application to Lennard-Jones cluster rearrangements.. 1998;108(22):9236–9245. doi:10.1063/1.476378.

10. Berryman JT, Schilling T. Sampling rare events in nonequilibrium and non-stationary systems. The Journal of Chemical Physics. 2010;133(24):244101. doi:10.1063/1.3525099.

11. Cérou F, Guyader A. Adaptive Multilevel Splitting for Rare Event Analysis. Stochastic Analysis and Applications. 2007;25(2):417–443. doi:10.1080/07362990601139628.

12. Allen RJ, Frenkel D, ten Wolde PR. Simulating rare events in equilibrium or nonequilibrium stochastic systems. The Journal of Chemical Physics. 2006;124(2):024102. doi:10.1063/1.2140273.

13. Russo JD, Zhang S, Leung JMG, Bogetti AT, Thompson JP, DeGrave AJ, et al. WESTPA 2.0: High-Performance Upgrades for Weighted Ensemble Simulations and Analysis of Longer-Timescale Applications. Journal of Chemical Theory and Computation. 2022;18(2):638–649. doi:10.1021/acs.jctc.1c01154.

14. Adhikari U, Mostofian B, Copperman J, Subramanian SR, Petersen AA, Zuckerman DM. Computational Estimation of Microsecond to Second Atomistic Folding Times. Journal of the American Chemical Society. 2019;141(16):6519–6526. doi:10.1021/jacs.8b10735.

15. Zwier MC, Pratt AJ, Adelman JL, Kaus JW, Zuckerman DM, Chong LT. Efficient Atomistic Simulation of Pathways and Calculation of Rate Constants for a Protein-Peptide Binding Process: Application to the MDM2 Protein and an Intrinsically Disordered p53 Peptide. The Journal of Physical Chemistry Letters. 2016;7(17):3440–3445. doi:10.1021/acs.jpclett.6b01502.

16. Lotz SD, Dickson A. Unbiased Molecular Dynamics of 11 min Timescale Drug Unbinding Reveals Transition State Stabilizing Interactions. Journal of the American Chemical Society. 2018;140(2):618–628. doi:10.1021/jacs.7b08572.

17. Saglam AS, Chong LT. Protein-protein binding pathways and calculations of rate constants using fully-continuous, explicit-solvent simulations. Chem Sci. 2019;10:2360–2372. doi:10.1039/C8SC04811H.

18. Sztain T, Ahn SH, Bogetti AT, Casalino L, Goldsmith JA, Seitz E, et al. A glycan gate controls opening of the SARS-CoV-2 spike protein. Nature Chemistry. 2021;13(10):963–968. doi:10.1038/s41557-021-00758-3.

19. Gillespie DT. Exact stochastic simulation of coupled chemical reactions. The journal of physical chemistry. 1977;81(25):2340–2361.

20. Zhang BW, Jasnow D, Zuckerman DM. The “weighted ensemble” path sampling method is statistically exact for a broad class of stochastic processes and binning procedures. The Journal of Chemical Physics. 2010;132(5):054107. doi:10.1063/1.3306345.

21. Reuter B, Fackeldey K, Weber M. Generalized Markov modeling of nonreversible molecular kinetics. The Journal of Chemical Physics. 2019;150(17):174103. doi:10.1063/1.5064530.

22. Reuter B, Klein M, Lange M. pyGPCCA - python GPCCA: Generalized Perron Cluster Cluster Analysis package to coarse-grain reversible and non-reversible Markov state models. 2022;doi:10.5281/zenodo.6914001.

23. Weber M, Galliat T. Characterization of Transition States in Conformational Dynamics using Fuzzy Sets. Takustr. 7, 14195 Berlin: ZIB; 2002. 02–12.

24. Deuflhard P, Weber M. Robust Perron cluster analysis in conformation dynamics. Linear Algebra and its Applications. 2005;398:161–184. doi:https://doi.org/10.1016/j.laa.2004.10.026.

25. Brandes U, Eiglsperger M, Lerner J, Pich C. Handbook of Graph Drawing and Visualization. Tamassia R, editor. Chapman and Hall/CRC.; 2013.

26. Bastian M, Heymann S, Jacomy M. Gephi: An Open Source Software for Exploring and Manipulating Networks. Proceedings of the International AAAI Conference on Web and Social Media. 2009;3(1):361–362.

27. Tse MJ, Chu BK, Gallivan CP, Read EL. Rare-event sampling of epigenetic landscapes and phenotype transitions. PLoS Comput Biol. 2018;14(8):e1006336. doi:10.1371/journal.pcbi.1006336.

28. Keating SM, Waltemath D, König M, Zhang F, Dräger A, Chaouiya C, et al. SBML Level 3: an extensible format for the exchange and reuse of biological models. Molecular systems biology. 2020;16(8):e9110.

